# The development of receptive field tuning properties in mouse binocular primary visual cortex

**DOI:** 10.1101/2022.02.14.480401

**Authors:** Liming Tan, Dario L. Ringach, Joshua T. Trachtenberg

**Author notes:** Correspondence: Joshua T. Trachtenberg.

## Abstract

The mouse primary visual cortex is a model system for understanding the relationship between cortical structure, function, and behavior (Seabrook et al., 2017; Chaplin and Margrie, 2020; Hooks and Chen, 2020; Saleem, 2020; Flossmann and Rochefort, 2021). Binocular neurons in V1 are the cellular basis of binocular vision, which is required for predation (Scholl et al., 2013; Hoy et al., 2016; La Chioma et al., 2020; Berson, 2021; Johnson et al., 2021). The normal development of binocular responses, however, has not been systematically measured. Here, we measure tuning properties of neurons to either eye in awake mice of either sex from eye-opening to the closure of the critical period. At eye-opening, we find an adult-like fraction of neurons responding to the contralateral-eye stimulation, which are selective for orientation and spatial frequency; few neurons respond to ipsilateral eye and their tuning is immature. Fraction of ipsilateral-eye responses increases rapidly in the first few days after eye opening and more slowly thereafter, reaching adult levels by critical period closure. Tuning of these responses improves with a similar time course. The development and tuning of binocular responses parallels that of ipsilateral-eye responses. Four days after eye-opening, monocular neurons respond to a full range of orientations but become more biased to cardinal orientations. Binocular responses, by contrast, lose their cardinal bias with age. Together, these data provide an in-depth accounting of the development of monocular and binocular responses in the binocular region of mouse V1 using a consistent set of visual stimuli and measurements.

**Significance statement:** In this manuscript, we present a full accounting of the emergence and refinement of monocular and binocular receptive field tuning properties of thousands of pyramidal neurons in mouse primary visual cortex. Our data reveal new features of monocular and binocular development that revise current models on the emergence of cortical binocularity. Given the recent interest in visually guided behaviors in mice that require binocular vision, e.g. predation, our measures will provide the basis for studies on the emergence of the neural circuitry guiding these behaviors.

## Introduction

Depth perception is central to many behaviors, including foraging and predation, and is thought to be a major driver of cortical evolution and brain size (Barton, 2004; Heesy, 2008). Binocular neurons are the neural substrate of depth perception. To support stereoscopic vision, binocular neurons must integrate inputs from the ipsilateral and contralateral eyes and these inputs must share similar receptive field tuning properties, including orientation, spatial frequency, and linearity (Bishop and Pettigrew, 1986; Wang et al., 2010; Tan et al., 2020). The normal development of binocular neurons, however, is incompletely understood.

Mice are a model system for studying visual cortical processing including binocular vision. Binocular neurons in mouse visual cortex encode disparity (Scholl et al., 2013, 2017a; La Chioma et al., 2019, 2020) and use this information to guide predation (Hoy et al., 2016; Samonds et al., 2019; Berson, 2021; Boone et al., 2021; Johnson et al., 2021). Characterizing the development of cortical responsiveness to vision through the contralateral and ipsilateral eyes and their integration to form binocular neurons would provide a foundation for understanding the molecular and cellular mechanisms that support the emergence of high acuity stereoscopic vision and visually guided behaviors. Here, we systematically measure the development of receptive field structure of monocular and binocular pyramidal neurons in layer 2/3 in awake, head-restrained mice over the first 3 weeks of normal vision from eye opening to postnatal day 36 (P36). We report a progressive increase in the fraction of binocular cells with age. Soon after eye opening, receptive field tuning of binocular neurons is less selective than the tuning of monocular neurons, and binocular neurons are largely responsive to stimuli oriented on the cardinal axes whereas monocular neurons respond to the obliques as well. By P36 binocular neurons are more selective than monocular neurons and have lost their cardinal bias. Binocular neurons at eye opening have poorly matched receptive field tuning for each eye, but this mismatch dissipates within 4 days after eye opening. Monocular neurons, by contrast, become more biased to cardinally oriented stimuli. Thus, there are progressive improvements in all aspects of binocular receptive field tuning that parallel similar decrements in monocular tuning. Visually evoked responses of binocular neurons, therefore, are preferentially improved in the weeks after eye opening.

## Materials and Methods

### Lead Contact

Further information and requests for resources and reagents should be directed to and will be fulfilled by the Lead Contact, Joshua Trachtenberg.

### Materials Availability

This study did not generate new unique reagents.

### Data and Code Availability

Custom-written MATLAB code and data for this study are available from the Lead Contact upon reasonable request.

### EXPERIMENTAL MODEL AND SUBJECT DETAILS

All procedures were approved by UCLA’s Office of Animal Research Oversight (the Institutional Animal Care and Use Committee, IACUC) and were in accord with guidelines set by the US National Institutes of Health. Mice were housed in groups of 2-3 per cage in a normal 12/12 light dark cycle. Animals were naïve subjects with no prior history of participation in research studies. A total of 25 mice, both male (M, n=18) and female (F, n=7) were used in this study (P14 layer 2/3, 8 M and 5 F; P18 layer 2/3, 4 M; P22, P29 and P36 layer 2/3, 4 M; P22, P29 and P36 layer 4, 2 M and 2 F).

Mice: All imaging was performed on mice expressing the slow variant of GCaMP6 in pyramidal neurons. For layer 2/3 imaging, these mice were derived from crosses of B6;DBA-Tg(tetO-GCaMP6s)2Niell/J (JAX Stock No: 024742; Wekselblatt et al., 2016) (28) with B6;CBA-Tg(Camk2a-tTA)1Mmay/J (JAX Stock No: 003010; Mayford et al., 1996) (29). For layer 4 imaging, these mice were derived from crosses of B6;C3-Tg(Scnn1a-cre)3Aibs/J (JAX Stock No: 009613) (Madisen et al., 2010) with Ai163 (Daigle et al., 2018) (Gift from Dr. Hongkui Zeng in Allen Institute). Mice expressing both transgenes were identified by PCR, outsourced to Transnetyx (transnetyx.com).

## METHODS DETAILS

### Surgery

All imaging experiments were performed through chronically-implanted cranial windows (Tan et al., 2020, 2021). In brief, mice were administered with carprofen prior to surgery, anesthetized with isoflurane (5% for induction; 1.5–2% during surgery), mounted on a stereotaxic surgical stage via ear bars and a mouth bar. Body temperature was maintained at 37°C via a heating pad. The scalp was removed, and the exposed skull was allowed to dry. The exposed skull and wound margins were then covered by a thin layer of Vetbond, followed by a thin layer of dental acrylic. A metal head bar was affixed with dental acrylic caudally to V1. A 3mm circular piece of skull overlying binocular V1 on the left hemisphere was removed using high-speed dental drill. A sterile 2.5 mm diameter cover glass was placed directly on the exposed dura and sealed to the surrounding skull with Vetbond. The remainder of the skull and the margins of the coverglass were sealed with dental acrylic. Mice were then recovered on a heating pad. When alert, they were placed back in their home cage. Carprofen was administered daily for 3 days post-surgery. Mice were left to recover for at least 3 days prior to imaging.

### Mapping of binocular area of the primary visual cortex

The location of binocular primary visual cortex for each mouse was identified using low magnification, epifluorescence imaging of GCaMP6s signals. Briefly, GCaMP6s was excited using a 470nm light-emitting diode. A 27-inch LCD monitor (ASUS, refreshed at 60 Hz) was positioned such that the binocular visual field fell in its center. The screen size was 112 deg in azimuth and 63 deg in elevation. The monitor was placed 20 cm from the eyes. A contrast reversing checkerboard (checker size 10×10 degree) bar windowed by a 1D Gaussian were presented along the horizontal or vertical axis to both eyes (Figure 1A). The checkerboard bar drifted normal to its orientation and swept the full screen width in 10 sec. Both directions of motion were used to obtain an absolute phase map along the two axes. Eight cycles were recorded for each of the four cardinal directions. Images were acquired at 10 frames per second with a PCO edge 4.2 sCMOS camera using a 35mm fixed focal length lens (Edmund optics, 35mm/F1.65, #85362, 3mm field of view). The visual areas were obtained from retinotopic maps of azimuth and elevation. The binocular area of the primary cortex was defined as the region of primary visual cortex adjacent to the higher visual area LM (Figure 1A).

**Figure 1:**
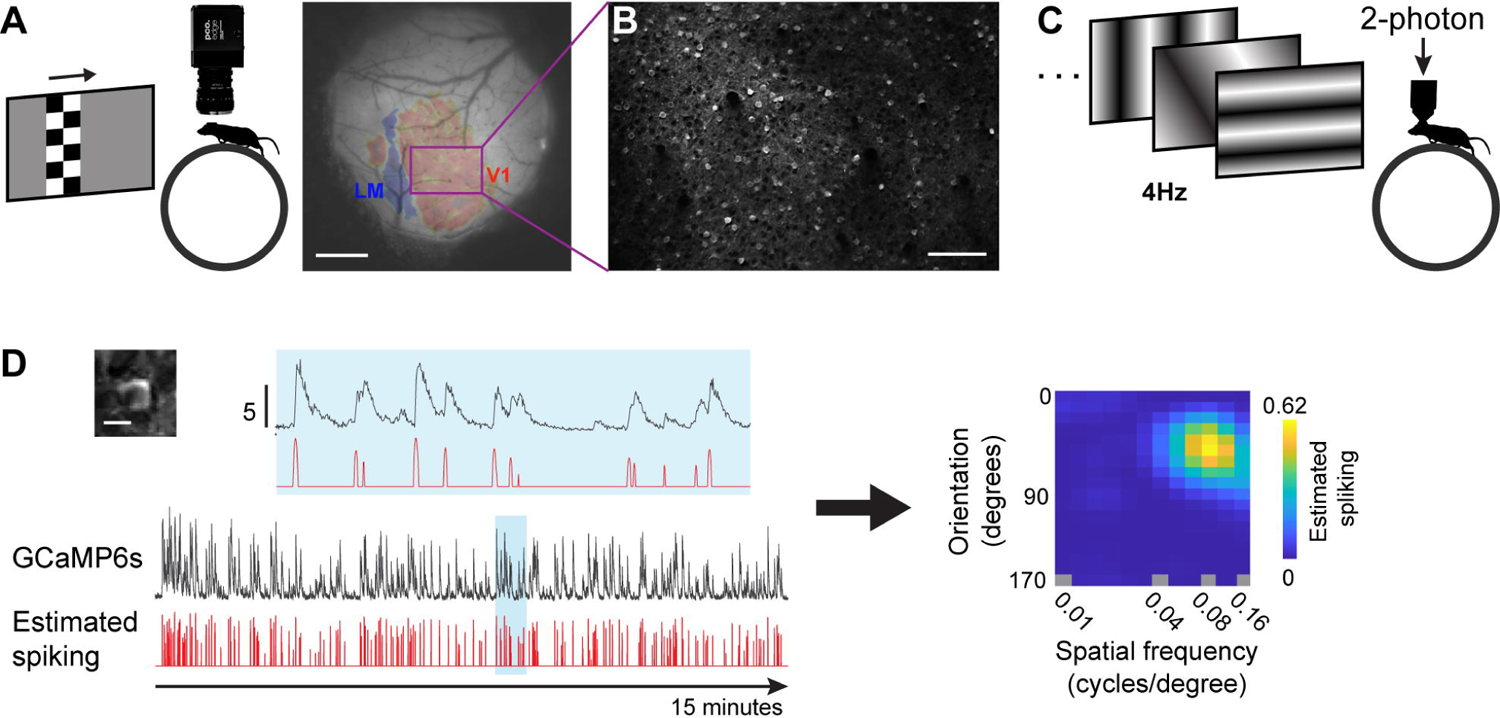
Overview of imaging and receptive field tuning A. Left: Mapping visual cortical areas using low-magnification epifluorescence imaging of GCaMP6s evoked responses to checkerboard bars, which were both drifting and flashing. Right: Example image of a cranial window highlighting the binocular primary visual cortex and the border with a higher visual area LM. The purple rectangle delineates the field of view used for 2-photon imaging. Scale bar, 0.5 mm. B. Field of view of in vivo 2-photon imaging of GCaMP6s labeled pyramidal neurons in the purple rectangle in panel A. Scale bar, 100μ C. Schematic of 2-photon imaging to a series of sinusoidal gratings sequentially presented at 4 Hz. Left: An image of a single neuron expressing GCaMP6s in panel B (Scale bar, 10μm). Below the image is the raw (black) and temporally deconvolved (red) GCaMP6s signal for 15 minutes of visual stimulation. The region in blue is expanded above horizontally to show the signal with greater detail. Scale bar, 5 dF/F0. Right: Receptive field tuning kernel of the cell on the left. Y-axis plots response strength as a function of stimulus orientation. X-axis plots response strength as a function of stimulus spatial frequency (on a log scale).

### Two-photon calcium imaging and visual stimulation

Two-photon imaging was done in the binocular area of V1 using a resonant/galvo scanning two-photon microscope (Neurolabware, Los Angeles, CA) controlled by Scanbox image acquisition software (Los Angeles, CA). A Coherent Discovery TPC laser (Santa Clara, CA) running at 920 nm focused through a 16x water-immersion objective lens (Nikon, 0.8 numerical aperture) was used to excite GCaMP6s. The objective was set at an angle of 10-11 degrees from the plumb line to reduce the slope of the imaging planes. Image sequences (512×796 pixels, 490×630μm, Figure 1B) were captured at 15.5 Hz at a depth of 120 to 300 μm below the pial surface on alert, head-fixed mice that were free to run on a 3D-printed running wheel (14cm diameter). A rotary encoder was used to record the rotations of this running wheel. To measure responses of neurons to each eye separately, an opaque patch was placed immediately in front of one eye when recording neuronal responses to visual stimuli presented to the other eye. On the screen that was used for visual area mapping, a set of static sinusoidal gratings were presented at 4 Hz in full screen in pseudo-random sequence with 100% contrast (Figure 1C). These gratings were generated in real-time by a Processing sketch using OpenGL shaders (see https://processing.org). These gratings are combinations of 18 orientations (equal intervals of 10° from 0° to 170°), 12 spatial frequencies (equal steps on a logarithmic scale from 0.0079 to 0.1549 cycles per degree) and 8 spatial phases. Imaging sessions were 15 min long (3600 stimuli in total), thus each combination of orientation and spatial frequency appeared 16 or 17 times.

Each of the 8 spatial phases for an orientation/spatial frequency combination appeared twice (F1/F0 values were calculated using responses of neurons as a function of spatial phase, see below). Transistor-transistor logic (TTL) signals were used to synchronize visual stimulation and imaging data. The stimulus computer generated these signals, which were sampled by the microscope electronics and time-stamped by the acquisition computer to indicate the frame and line number being scanned at the time of the TTL.

### Analysis of two-photon imaging data

#### Image processing

The pipeline for image processing has been described in detail (Tan et al., 2020, 2021). Briefly, movies from the same plane for each eye were concatenated and motion corrected. Regions of interest (ROI) corresponding to pyramidal neuron soma were determined using a Matlab graphical user interface tool (Scanbox, Los Angeles, CA).

Using this GUI, we computed pixel-wise correlations of fluorescence changes over time. The temporal correlation of pixels was used to determine the boundary of ROI for each neuron. After segmentation, the fluorescence signal for each ROI and surrounding neuropil was extracted. The neuropil signal for a given ROI was computed by dilating the ROI with a disk of 8-pixel radius. The original ROI and those of other cells that overlap with the region were excluded, and the average signal within this area was computed. The signal obtained from the ROI was then robustly regressed on the neuropil. The residual represented the corrected signal of the ROI. The correction factor was derived from the slope of the robust regression. Neuronal spiking was estimated via non-negative temporal deconvolution of the corrected ROI signal using Vanilla algorithm (Berens et al., 2018). Subsequently, fluorescent signals and estimated spiking for each cell were split into separate files corresponding to the individual imaging session for each eye.

### Calculation of response properties

#### Identification of visually responsive neurons using SNR

Signal to noise ratio (SNR) was used to identify neurons with significant visual responses. SNR for each neuron was calculated based on the optimal delay of the neuron. Optimal delay was defined as the imaging frame after stimulus onset at which the neuron’s inferred spiking reached maximum. To calculate SNR, signal was the mean of standard deviations of spiking to all visual stimuli at the optimal delay (5-7 frames, thus ∼0.387 sec, after stimulus onset), and noise was this value at frames well before or after stimulus onset (frames –2 to 0, and 13 to 17). Neurons whose optimal delays occurred outside of the time-locked stimulus response window of 4 to 8 frames (padded by ±1 frame around the 5-7 frame range used above) after stimulus onset were spontaneously active but visually unresponsive. They had SNR values close to 1. The SNR values of these unresponsive neurons were normally distributed (mean=1.0) over a narrow range. Unresponsive neurons with optimal delays naturally occurring in the 4-8 frame time window can be distinguished from visually responsive neurons by SNR. This SNR threshold was defined at 3 standard deviations above the mean SNR of the above-mentioned normal distribution. SNR values were calculated separately for responses to the ipsilateral or contralateral eye. Visually evoked responses of neurons had optimal delays between frames 4 and 8, and SNRs greater than this cutoff. If responses evoked via stimulation of each eye, separately, met these criteria, the cell was considered to be binocular.

#### Tuning kernel for orientation and spatial frequency

The estimation of the tuning kernel was performed by fitting a linear model between the response and the stimulus (Ringach et al., 2016). Cross-correlation maps were used to show each neuron’s spiking level to visual stimuli (orientation and spatial frequency) by averaging responses over spatial phases. The final tuning kernel of a neuron was defined as the correlation map at the optimal delay (Figure 1D). An advantage of reverse correlation methods is that they tend to linearize the neurons around their operating point (Ringach and Shapley, 2004). Thus, if the tuning of the cell can be modeled as a linear kernel followed by a static nonlinearity (either compressive or expansive), the recovered kernel will be a faithful representation of its tuning up to a multiplicative constant (Ringach et al., 1997b; Chichilnisky, 2001). This makes it possible to compare tuning properties across neurons and over time.

#### F1/F0 measurement for response linearity

F1/F0 is the ratio of the 1^st^ Fourier harmonic and 0^th^ Fourier harmonic for a given neuron across different spatial phases (Ringach et al., 2002). For complex cells the F1/F0<1, while for simple cells F1/F0>1(Skottun et al., 1991).

#### Orientation and spatial frequency preference

We used horizontal (for spatial frequency) and vertical (for orientation) slices of the tuning kernel through the peak inferred spiking to calculate orientation and spatial frequency preferences.

Orientation preference:

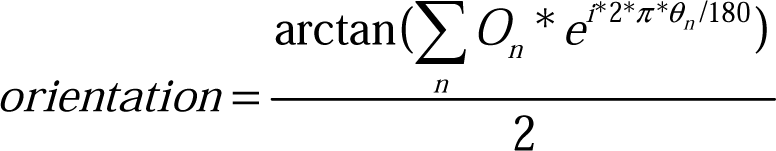

O_n_ is a 1×18 array, in which a level of estimated spiking (O_1_ to O_18_) occurs at orientations θ_n_ (0° to 170°, spaced every 10°). Orientation is calculated in radians and then converted to degrees.

Spatial frequency preference:

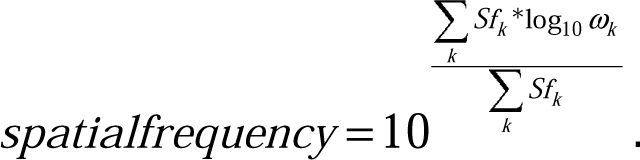

Sf_k_ is a 1×12 array, in which a level of estimated spiking (Sf_1_ to Sf_12_) occurs at spatial frequencies ω_k_ (12 equal steps on a logarithmic scale from 0.0079 to 0.1549 cycles per degree).

#### Circular variance

Circular variance is a measure of orientation selectivity, ranging from zero (highest selectivity) to one (lowest selectivity). The circular variance of a neuron whose estimated spiking, O_n_, occurred at orientations θ_n_ (0° to 170°, spaced every 10°), is defined as

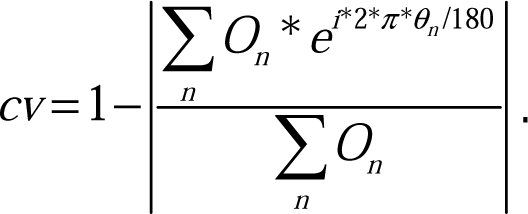

#### Cardinal proportion

Polar histograms showing orientation distributions (Figures 3A and 7D) were used to calculate cardinal proportion of monocular and binocular neurons. These polar histograms were plotted using bins of 11.25° (16 bins across 180°). In this study, cardinal orientation preference was defined within 0°±16.875° (degrees represented by 1.5 bins, 11.25°x1.5) or 90°±16.875°. Proportions of neurons with cardinal orientation preference can then be defined as

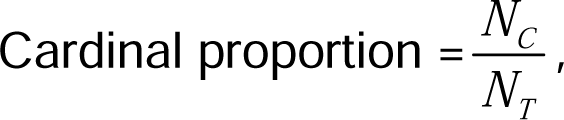

where *N_C_* is the number of monocular or binocular neurons whose orientation preferences were within the cardinal orientation range, *N_T_* is total number of neurons. For binocular neurons, the plotted cardinal proportion is the mean of cardinal proportions for contralateral and ipsilateral eye responses (Figures 3C, 4C and 7F, G).

#### ΔOrientation for binocular neurons

For a binocular neuron, *Ori_contra_* and *Ori_ipsi_* are the neuron’s orientation preferences to contralateral and ipsilateral eye, respectively.

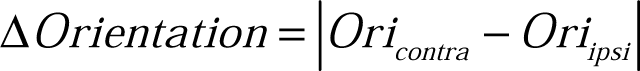

If the value of Δ*Orientation* is above 90 (e.g. |170-10|=160), then the actual value for the difference of orientation preferences to two eyes is 180-Δ*Orientation* (180-160=20).

#### Binocular matching coefficient

This was defined as the correlation coefficient between contralateral and ipsilateral tuning kernels of binocular neurons.

### Figure plotting

#### Density profile plots

The code for calculating density profiles was modified from Matlab code scattercloud (https://www.mathworks.com/matlabcentral/fileexchange/6037-scattercloud). Briefly, we first made same number of bins (n=11∼16) along both the x- and y-axis for measurements used in scatter plot. We then calculated density of data points in each bin to get an n-by-n density profile matrix and plot the matrix using Matlab surf function with interpolated coloring for each face.

#### Density profiles overlay

We overlaid pairs of density profiles by using Matlab imfuse function. Before overlaying two matrices, we normalized each matrix to limit density between 0 and 1 to make the two density profiles being merged in the same scale.

#### Quantification and Statistical Analysis

Sample size was not determined by a-priori power analysis. All statistical analyses were performed in MATLAB (https://www.mathworks.com/) using non-parametric tests with significance levels set at α < 0.05, and did Bonferroni corrections on α for multiple comparisons when necessary. Mann-Whitney U-tests (Wilcoxon rank sum test) were used to test differences between two independent populations. When comparing more than two populations that were non-normally distributed, a Kruskal-Wallis test, a nonparametric version of one-way ANOVA, was used to determine whether statistically significant differences existed among these independent populations. If significant differences did exist, post hoc multiple comparison tests or Mann-Whitney U-tests with Bonferroni corrections were used to test for significant differences between pairs within the group.

## Results

### Measuring receptive field tuning in binocular visual cortex

Binocular visual cortex was mapped via epifluorescence imaging and receptive field tuning was measured via 2-photon calcium imaging followed published protocol (Tan et al., 2020, 2021). In brief, for each mouse in this study, the binocular region was identified using retinotopic mapping of GCaMP6s responses (Figure 1A). These maps and the corresponding maps of vasculature were used to target high-resolution 2-photon calcium imaging of single neurons (Figure 1B). To measure receptive field tuning of excitatory neurons in layer 2/3 and 4, 2-photon imaging was used to record fluorescence changes in neurons expressing GCaMP6s in alert, head-fixed mice viewing a battery of flashed sinusoid gratings comprising 18 orientations of 12 spatial frequencies and 8 spatial phases presented at 4Hz (Figure 1C). For each imaged neuron, its receptive field tuning was estimated from the linear regression of the temporally deconvolved calcium response. The resultant “tuning kernel” plots response strength across orientations and spatial frequencies (Figure 1D).

### Development of monocular and binocular responses

Mice open their eyes on or around postnatal day 14 (P14). At eye opening, only about half of imaged neurons responded to our stimulus battery (Figure 2A, B, D). Of those, the vast majority responded only when stimulated via the contralateral eye (46% of all imaged neurons, 84% of visually responsive neurons; Figure 2C, E). Over the next 4 days after eye opening (P14-P18), the fraction of visually responsive neurons increased to about three-quarters of imaged neurons. This was largely due to neurons becoming responsive to stimulation of the ipsilateral eye. The gain of cortical responsiveness to ipsilateral eye stimulation also drove the expansion of the binocular pool. During this period, the fraction of neurons responding solely to contralateral eye stimulation decreased significantly (Figure 2C, E). Earlier work indicates that this does not result from a large loss of contralateral eye responsiveness per-se, but from the conversion of many monocular neurons to binocular (Tan et al., 2021). That is, many neurons that were previously responsive solely to the contralateral eye gained responsiveness to the ipsilateral eye and became binocular. Thus, the fraction of neurons responding to contralateral eye stimulation, whether it be monocular, or binocular contracted only slightly (the sum of the green and blue points in Figure 2C, E) even as the purely monocular fraction dropped considerably.

**Figure 2:**
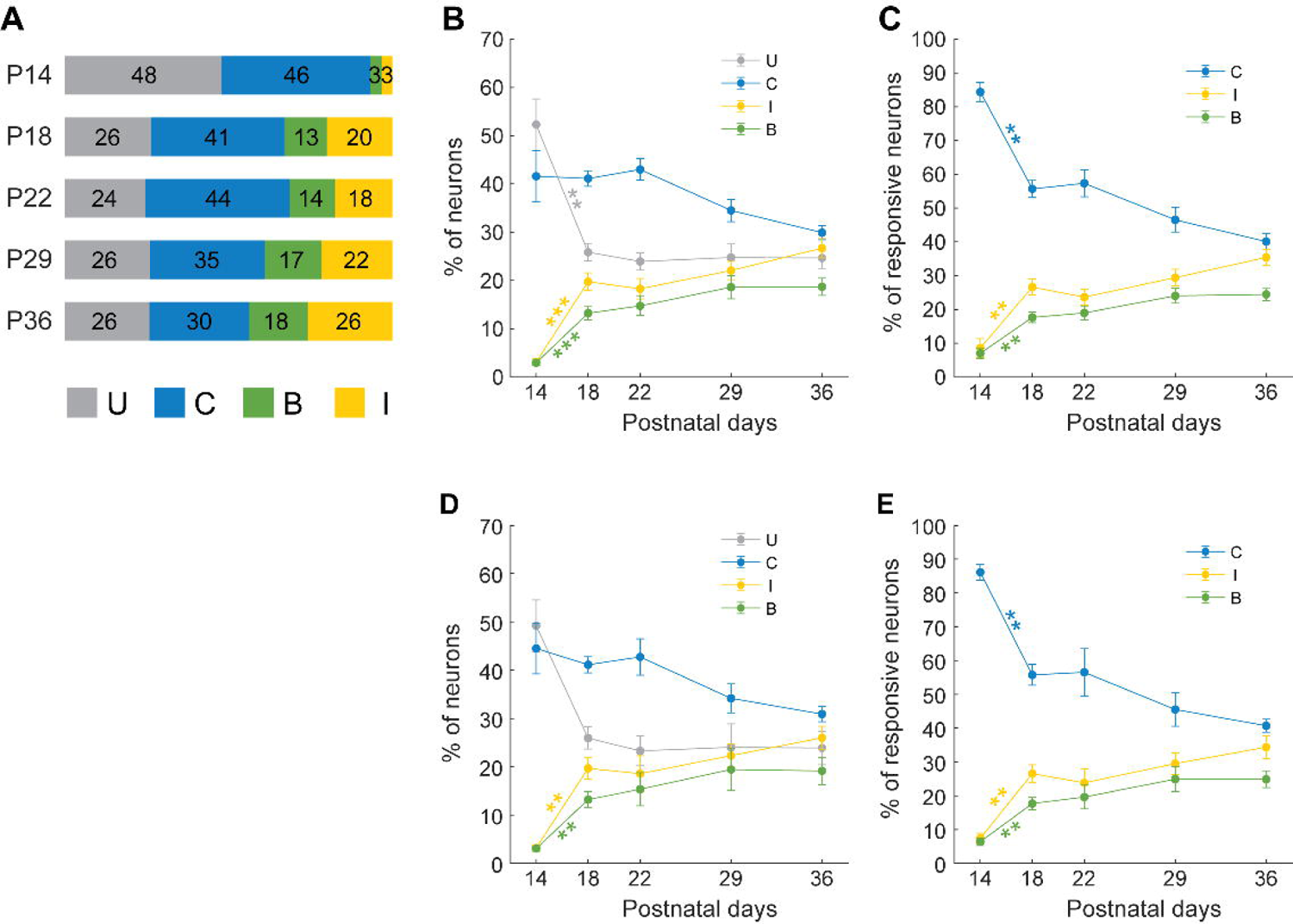
Fraction of visually responsive neurons in layer 2/3 at each age A. Proportions of all imaged neurons in layer 2/3 at each age that are unresponsive to our visual stimuli (gray), that respond solely to stimulation of the contralateral eye (blue), respond to stimulation of either eye (green), or respond solely to stimulation of the ipsilateral eye (yellow). P14, 2227 cells, 13 mice; P18, 1708 cells, 4 mice; P22, 3268 cells, 4 mice; P29, 2389 cells, 4 mice; P36, 1905 cells, 4 mice. B. Proportions of neurons per field of view (FOV) as a function of age in V1B layer 2/3. P14, 15 FOV; P18, 8 FOV; P22, 11 FOV; P29, 11 FOV; P36, 11 FOV. Mean and standard error of the mean (SEM) at each time point were shown as dots and error bars. Statistics: Mann-Whitney U tests with Bonferroni corrections on data between adjacent days. **, p<0.01; ***, p<0.001. See also Figure 2-1. C. As in B but only for visually responsive neurons. See also Figure 2-1. D. Proportions of neurons per mouse as a function of age in V1B layer 2/3. P14, 13 mice; P18, 4 mice; P22, 4 mice; P29, 4 mice; P36, 4 mice. Mean and standard error of the mean (SEM) at each time point were shown as dots and error bars. Statistics: Mann-Whitney U tests with Bonferroni corrections on data between adjacent days. **, p<0.01. See also Figure 2-1. E. As in D but only for visually responsive neurons. See also Figure 2-1.

From P18 to P36, these fractions shifted more modestly (Figure 2C, E). Over these weeks, there was a progressive reduction in the fraction of cells that responded solely to contralateral eye stimulation, and this reduction was paralleled by gradual increases in the fractions of cells that responded when stimulated via either eye or only via the ipsilateral eye. By P36, about 40% of visually responsive neurons were monocularly responsive to the contralateral eye, roughly 35% were monocularly responsive to the ipsilateral eye, and another 25% were binocular. This high degree of monocularity in the binocular zone is in agreement with others (Salinas et al., 2017).

### Orientation preference and binocular matching

At eye opening, neurons driven solely by stimulation of the contralateral eye showed a strong bias to vertically oriented sinusoid gratings (Figure 3A-C; blue). By contrast, the few neurons that were responsive solely to the ipsilateral eye responded to a broader range of orientations at eye opening (Figure 3A-C; yellow). Within the first 4 days after eye opening, this bias was lost and monocular, contralateral eye-evoked responses could be driven by a range of orientations. This broadening of orientation representation was, however, short lived. By P22 and remaining through P36, monocular, contralateral neurons exhibited a bias to the cardinal axes. Similarly, monocular, ipsilateral eye-evoked responses showed little orientation bias at P18 but gradually gained a cardinal bias from P18 and P36. Thus, by P36, similar cardinal biases were seen for monocular ipsilateral and monocular contralateral neurons. These measures are in agreement with previously published data reporting a strong cardinal bias in monocular neurons in mouse primary visual cortex (Roth et al., 2012; Salinas et al., 2017; Scholl et al., 2017b). Binocular neurons followed a different trajectory (Figure 3A-C; green). At P18, their responses were strongly biased towards the cardinal axes (mostly horizontal), and this bias was progressively lost with age. Thus, monocular neurons become progressively more biased to stimuli oriented on the cardinal axes while binocular neurons come to encode the full range of orientations.

**Figure 3.**
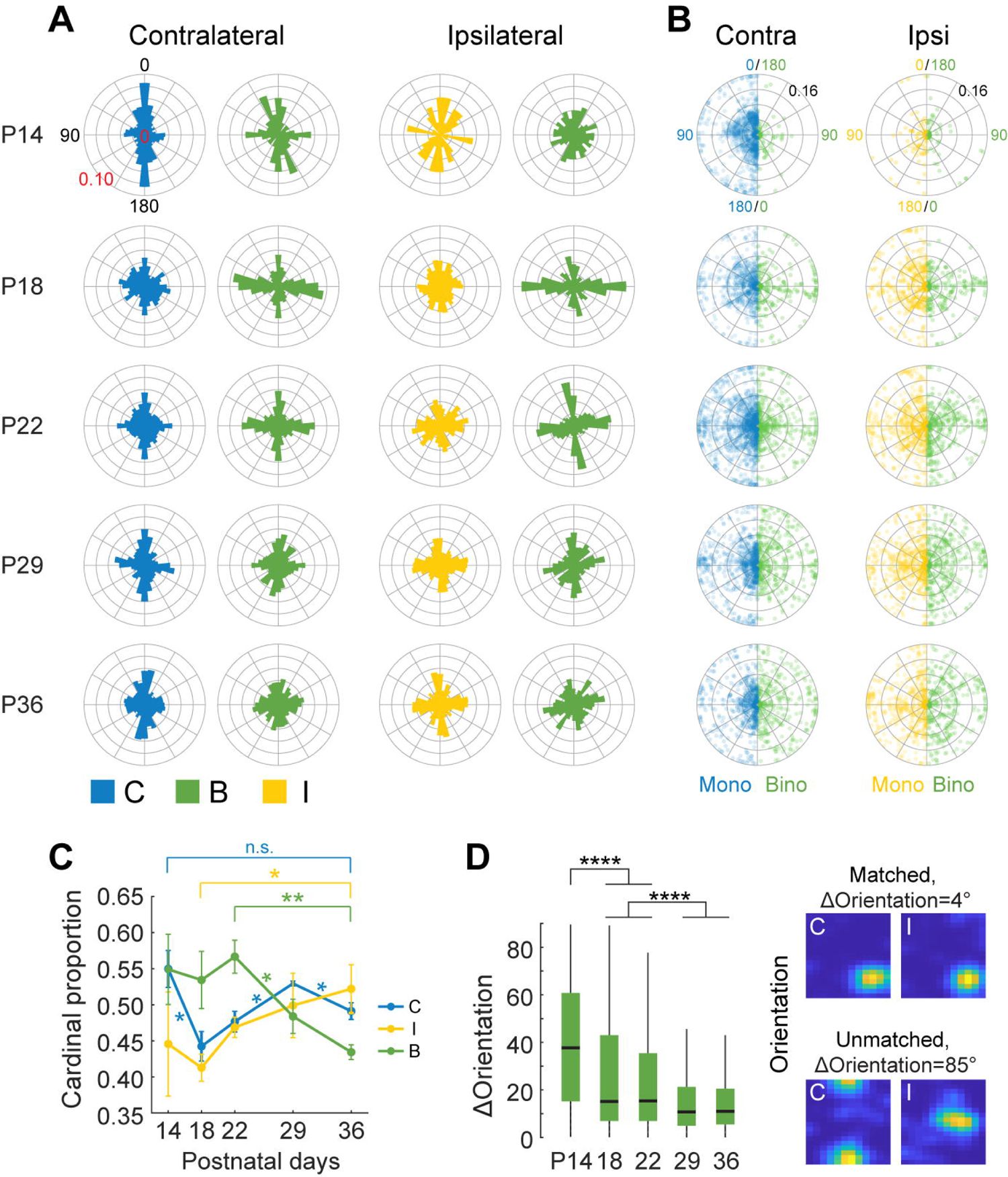
Orientation preference of monocular and binocular neurons in layer 2/3 as a function of age A. Polar histograms depicting the fractions of visually responsive neurons in layer 2/3 preferring to a particular orientation. Note that the plots are mirror symmetric from top to bottom. Stimuli were not drifting, and thus the orientations span 0 to 180 degrees. Plots are color coded to represent neurons that responded solely to contralateral (blue) or ipsilateral (yellow) eye stimulation, or to stimulation of either eye (binocular; green). For binocular neurons, the orientation preferences obtained via stimulation of the contralateral or ipsilateral eyes are plotted separately. Monocular contralateral neurons: P14, n=1015; P18, n=696; P22, n=1442; P29, n=834; P36, n=581. Monocular ipsilateral neurons: P14, n=73; P18, n=342; P22, n=574; P29, n=514; P36, n=493. Binocular neurons: P14, n=74; P18, n=220; P22, n=450; P29, n=417; P36, n=339. B. Polar scatter plots showing the orientation and spatial frequency preferences of all visually responsive layer 2/3 neurons. Each dot is a neuron. In each plot, monocular neurons were shown in the left half and binocular neurons were shown in the right half. Radius of plots represent spatial frequencies that spaced evenly from 0 to 0.16 cycles per degree. Dots were color coded as in A with ∼20% opacity to highlight dense regions. C. Proportions of neurons with cardinal orientation preferences per mouse. See methods for detail. Mean and standard error of the mean (SEM) at each time point were shown as dots and error bars. Two-sample t-test between adjacent days as well as between P14 and P36 for C, between P18 and P36 for I, and between P22 and P36 for B. *, p<0.05; **, p<0.01. Note from P18 to P36 the increase of cardinal proportion for monocular neurons, and decrease in cardinal proportion for binocular neurons. D. Left: boxplots of the differences in orientation preferences to either eye of binocular neurons. Kruskal-Wallis test followed by multiple comparison test with Bonferroni corrections, **** P<0.0001. Right: Tuning kernels to the contralateral (C) or ipsilateral eye (I) from a matched binocular neuron (top) and an unmatched binocular neuron (bottom) in layer 2/3. Kernels for each neuron were normalized to the peak inferred spiking of the neuron. The difference in orientation preference to one or the other eye, or Δthe kernels for each neuron.

Binocular neurons integrate visual information received from the two eyes. The eyes are spatially offset in the head and thus convey slightly different images to cortex. Depth perception only emerges when binocular neurons successfully integrate similar information from the two eyes. This integration is reflected in their receptive field tuning properties, which match for each eye (Wang et al., 2010; Sarnaik et al., 2014; Gu and Cang, 2016; Chang et al., 2020; Tan et al., 2020). Thus, if a neuron responds optimally to a particular orientation and spatial frequency stimulus presented to one eye, it will optimally respond to highly similar stimulus presented to the other. Any deviations in preferences between the two eyes is known as binocular mismatch. At eye opening, binocular neurons were largely mismatched in their orientation tuning preferences measured from each eye, with a difference of about 40+/-20 degrees (Figure 3D). By P18, near adult levels of binocular matching were achieved, though there were small but significant improvements through P36. While the median difference dropped only marginally over this time, the distributions tightened considerably (Figure 3D).

### Spatial frequency tuning and cardinality

Previous recordings from monocular neurons found a correlation between cardinal orientation bias and high spatial frequency preferences (Salinas et al., 2017). Specifically, neurons that responded optimally to spatial frequencies above 0.24 cycles per degree (cpd) tended to also respond optimally to cardinal orientations. Given this, we examined whether the progressive shift towards greater cardinality in monocular neurons, but towards less cardinality in binocular neurons could more simply be attributed to spatial frequency preference, rather than ocularity. We found a relationship between cardinality and spatial frequency in young mice (P14-P22), but this relationship was the same for monocular and binocular neurons (Figure 4A-C). Specifically, neurons responding optimally to spatial frequencies between 0.08cpd and 0.16cpd had greater cardinal biases than those that responded optimally to spatial frequencies below 0.08cpd. This relationship was less evident in older mice (Figure 4A-C). Thus, the developmental shifts in cardinality of monocular and binocular neurons are unlikely to be due to spatial frequency preference.

**Figure 4.**
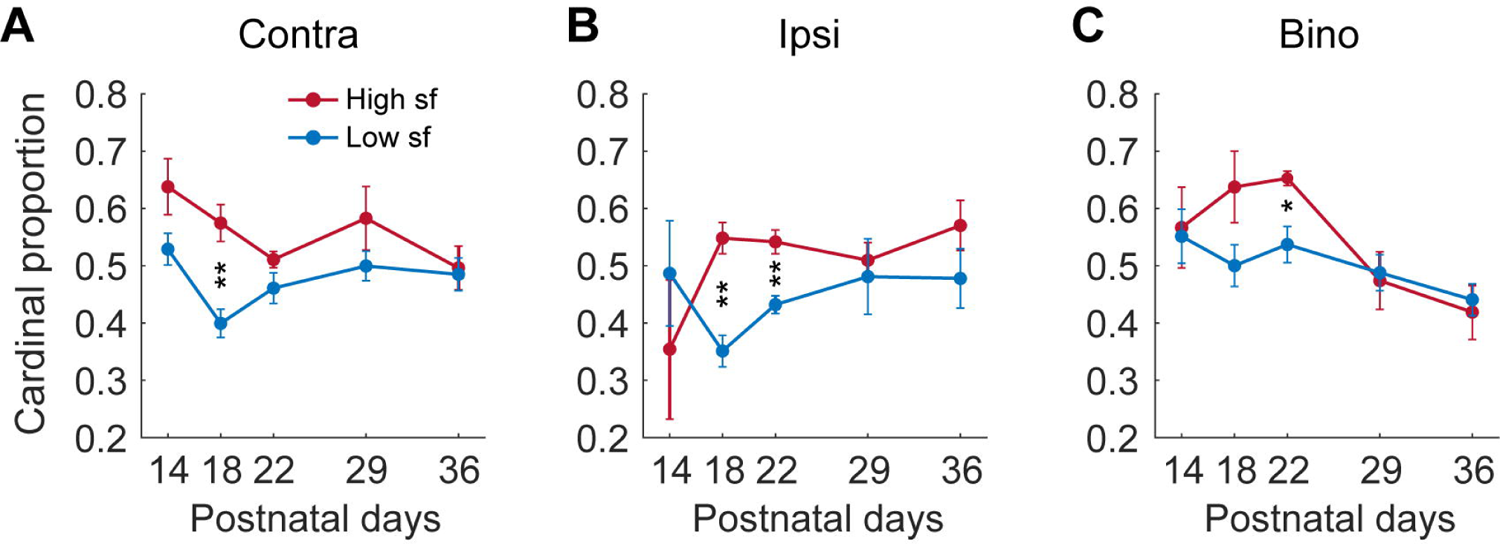
Relationship between spatial frequency tuning and cardinality during development. A. Cardinal proportions for monocular contralateral neurons per mouse preferring high (High sf, red, >=0.08cpd) or low spatial frequency (Low sf, blue, <0.08cpd), respectively. Mean and standard error of the mean (SEM) at each time point were shown as dots and error bars. Two-sample t-test between high and low sf preferring neurons on each day. *, p<0.05; **, p<0.01. B. As in A but for monocular ipsilateral neurons. C. As in A but for binocular neurons.

### Changes in receptive field tuning of monocular and binocular neurons

We also characterized the progression of orientation selectivity, spatial frequency preference, and linearity (Figure 5). Orientation selectivity is measured as circular variance (Ringach et al., 1997a). In this measure, smaller values indicate sharper tuning and greater selectivity. Linearity refers to response sensitivity to the spatial phase of the sinusoid. If a cell responds to sinusoid grating of only a particular spatial phase it is referred to as a “simple” cell; those whose responses are insensitive to the spatial phase of the sinusoid are “complex”. The ratio of the 1^st^ Fourier harmonic and 0^th^ Fourier harmonic across different spatial phases can be used to classify such responses (Skottun et al., 1991; Ringach et al., 1997a; Mechler and Ringach, 2002). Broadly, for complex cells F1/F0<1, while for simple cells F1/F0>1.

**Figure 5.**
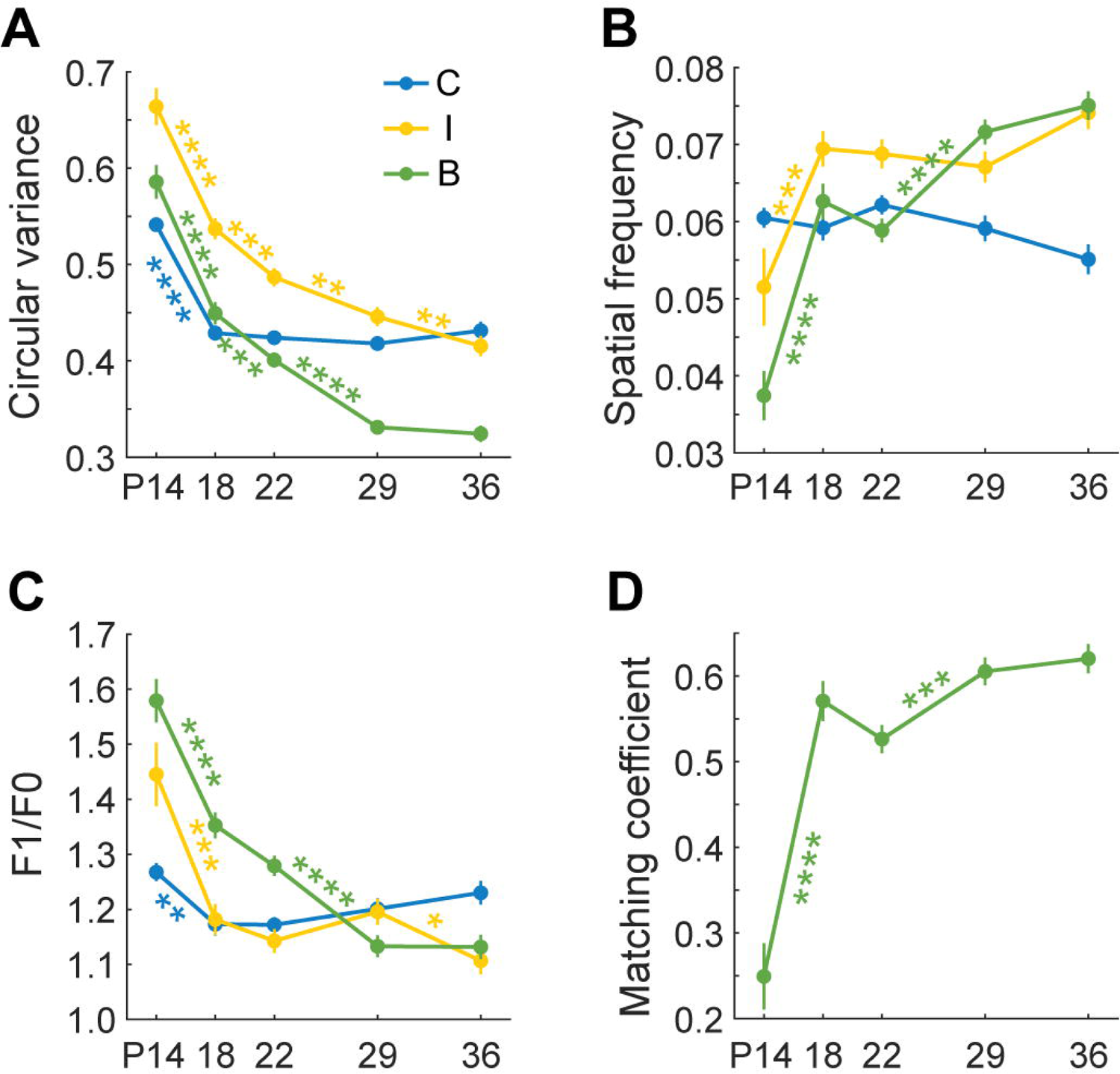
The progression of layer 2/3 receptive field tuning properties from eye opening to P36 A. Plots of the mean circular variance as a function of age for cells in layer 2/3 that are solely responsive to stimulation of the contralateral (blue) or ipsilateral (yellow) eye or are responsive to either eye (binocular; green). Error bars are standard error of the mean. Mann-Whitney U tests with Bonferroni corrections on data between adjacent days. *, p<0.05; **, p<0.01; ***, p<0.001; ****, p<0.0001. B. As in A, but for spatial frequency preference. C. As in A, but for response linearity (F1/F0). D. Plot of mean matching coefficients for binocular neurons as a function of age. Error bars are standard error of the mean. Mann-Whitney U tests with Bonferroni corrections on data between adjacent days.

At eye opening, monocular responses to contralateral eye stimulation, which constituted most evoked responses, were characterized by broad orientation selectivity (high circular variance) but adult-like levels of spatial frequency preferences and response linearities (Figure 5A-C, blue). By P18, 4 days after eye opening, monocular contralateral eye responses attained adult-like orientation selectivity. By comparison, the few neurons that responded to monocular ipsilateral eye stimulation at eye opening were less selective for orientation, preferred lower spatial frequencies and had more linear responses (Figure 5A-C, yellow). Ipsilateral-eye evoked orientation improved exponentially thereafter. The greatest improvements occurred within the first week after eye opening (P14-21), but further improvements in orientation selectivity continued through P36. Measures of spatial frequency preference and linearity improved quickly between P14 and P18 but were relatively stable thereafter.

The development of binocular receptive field tuning progressed from being quite poor at eye opening, relative to the tuning of monocular neurons, to becoming more selective than monocular neurons by P36. All tuning measures improved somewhat linearly and continuously from P14 through P36 (Figure 5A-C, green). We have addressed the mechanisms underlying this transition in earlier work (Tan et al., 2020, 2021) and these involve the progressive recruitment of the best tuned monocular neurons into the binocular pool as poorly tuned binocular neurons lose responsiveness to one eye and become monocular. Binocular matching coefficients, measured as the correlation coefficient of each eye’s tuning kernels, were quite poor at eye opening but experienced a large increase (better matching) by P18 with modest improvement thereafter (Figure 5D). In agreement with others, differences in binocular matching were most evident for orientation tuning and less so for spatial frequency (Salinas et al., 2017). All measures from all cells at each age are given in Figure 6.

**Figure 6.**
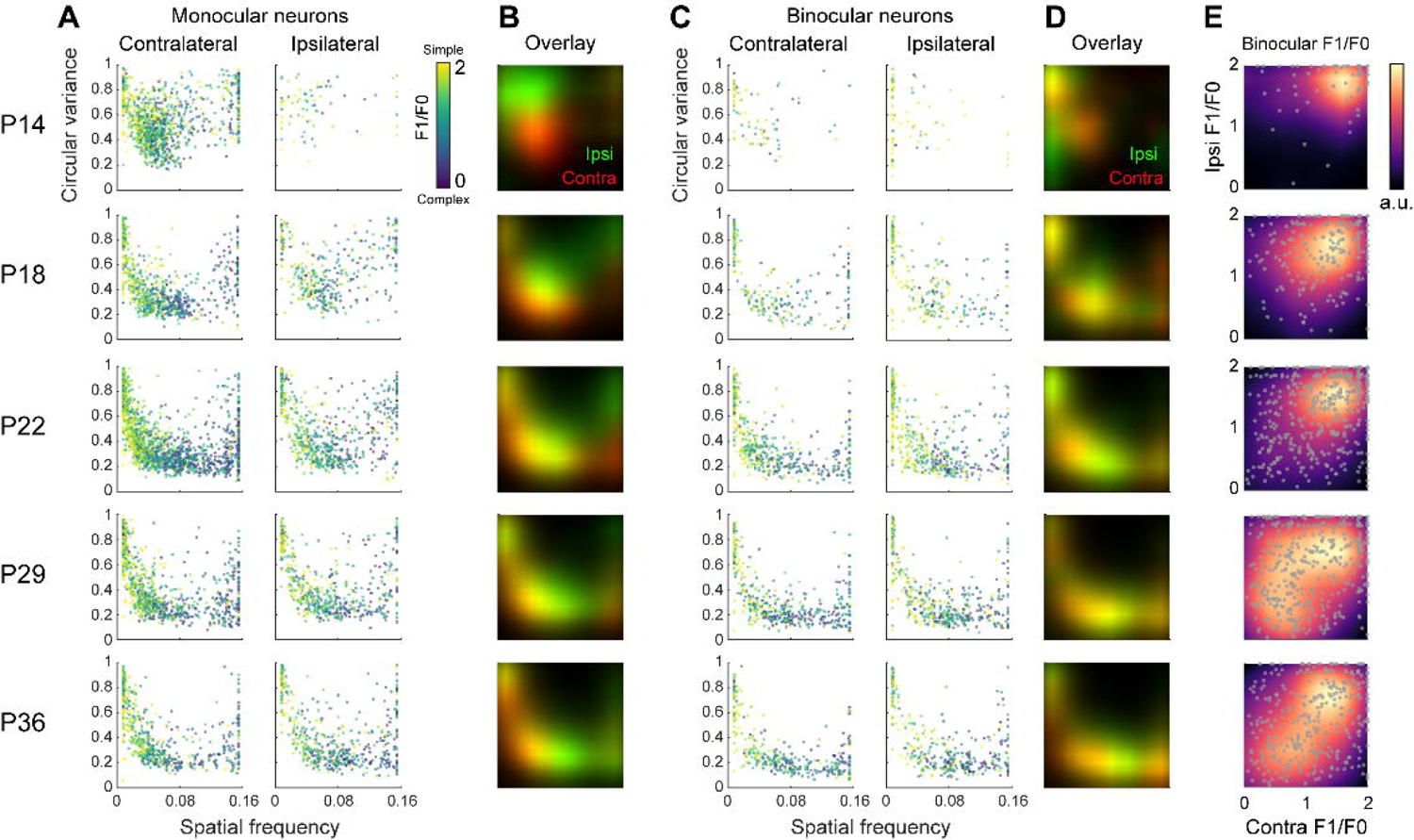
All data points for monocular and binocular neurons as a function of age A. Plots of the circular variance and spatial frequency preferences for all monocular neurons at each age. In each plot, the linearity (F1/F0) of the response is color coded as shown in the color bar to the right. Yellow is simpler, while dark blue is more complex. Left column shows tuning preferences for monocular, contralateral eye responsive neurons. Right column shows tuning preferences for monocular, ipsilateral eye responsive neurons. B. Overlaid tuning density profile maps showing the relative distribution of tuning of neurons responding solely to contralateral or ipsilateral eye stimulation. Note that ipsilateral eye responses are initially more poorly tuned, but progressively improve to match the tuning of contralateral eye evoked responses. Red: contralateral eye responses. Green: ipsilateral eye responses. C. As in A, but for binocular neurons. Response tuning obtained via stimulation of the contralateral and ipsilateral eyes were plotted separately. D. As in B, but for binocular neurons. E. Scatter plots on top of density profile maps showing the density distribution of response linearity obtained via stimulation of a binocular neuron through the contralateral eye (X-axis) and ipsilateral eye (Y-axis). Each point is a single cell. Note that responses are quite simple at eye opening and become progressively more complex with age.

### Ipsilateral eye responses are less selective in layer 4 than in layer 2/3

We could not measure the early development of visually evoked responses in layer 4. In the Scnn1a-Tg3-Cre mice that we used to restrict expression of GCaMP6s to layer 4, Cre is not expressed until ∼P21, about a week after eye opening. From our observations of receptive field tuning from P22 through P36, we found that responses in layer 4 become progressively more monocular with age (Figure 7A-C), as opposed to more binocular in layer 2/3. Proportions of neurons in layer 4 preferring cardinal orientations did not change from P22 to P36 (Figure 7D-G). Moreover, there was no significant improvement in binocular matching with age in layer 4 (Figure 7H). Orientation selectivity of contralateral-eye evoked responses was similar between layer 4 and 2/3, while ipsilateral-eye evoked responses were more broadly tuned in layer 4 (Figure 8A). Layer 4 neurons were typically responsive to higher spatial frequency stimuli than neurons in layer 2/3 but were more linear/simple (Figure 8B,C). Thus, there are distinct differences in receptive field tuning properties of neurons in layers 2/3 and 4, especially for ipsilateral-eye evoked responses or binocular responses which are significantly more sharply tuned in layer 2/3 than in layer 4.

**Figure 7.**
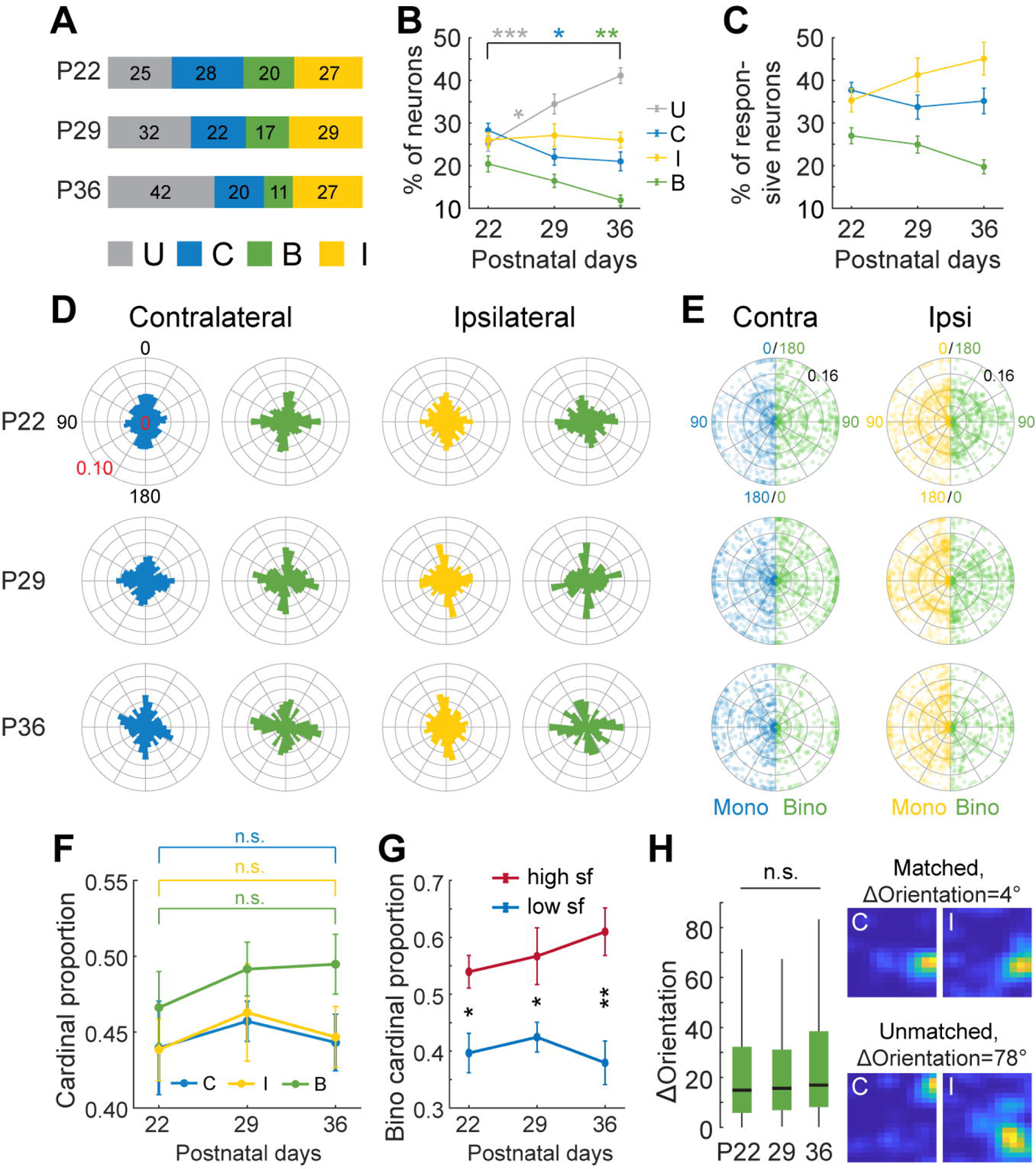
Responsiveness and orientation preferences in layer 4 A. Proportions of all imaged neurons in layer 4 at each age that are unresponsive to our visual stimuli (gray), that respond solely to stimulation of the contralateral eye (blue), respond to stimulation of either eye (green), or respond solely to stimulation of the ipsilateral eye (yellow). P22: 1391 cells, 4 mice; P29: 2526 cells, 4 mice; P36: 2190 cells, 4 mice. B. Proportions of neurons per FOV as a function of age in V1B layer 4. P22, 12 FOV; P29, 12 FOV; P36, 12 FOV. Mean and standard error of the mean (SEM) at each time point were shown as dots and error bars. Statistics: Mann-Whitney U tests with Bonferroni corrections on data between adjacent days, as well as between P22 and P36. *, p<0.05; **, p<0.01; ***, p<0.001. C. As in B but only for visually responsive neurons. D. Polar histograms as in Figure 3A showing the fraction of responsive neurons in layer 4 preferring to a particular orientation. Monocular contralateral neurons: P22, n=552; P29, n=546; P36, n=425. Monocular ipsilateral neurons: P22, n=528; P29, n=730; P36, n=600. Binocular neurons: P22, n=392; P29, n=428; P36, n=248. E. Polar scatter plots as in Figure 3B showing the orientation and spatial frequency preferences of all visually responsive layer 4 neurons. F. Proportions of L4 neurons with cardinal orientation preferences per mouse. Two-sample t-test between adjacent days as well as between P22 and P36. G. Cardinal proportions for binocular layer 4 neurons per mouse preferring high (high sf, red, >=0.08cpd) or low spatial frequency (low sf, blue, <0.08cpd), respectively. Mean and standard error of the mean (SEM) at each time point were shown as dots and error bars. Two-sample t-test between high and low sf preferring neurons on each day. *, p<0.05; **, p<0.01. H. Left: boxplots of the differences in orientation preferences to either eye of layer 4 binocular neurons. Kruskal-Wallis test followed by multiple comparison test with Bonferroni corrections. Right: Tuning kernels to the contralateral (C) or ipsilateral eye (I) from a matched binocular neuron (top) and an unmatched binocular neuron (bottom) in layer 4. Kernels for each neuron were normalized to the peak inferred spiking of the neuron. ΔOrientation was shown above the kernels for each neuron.

**Figure 8.**
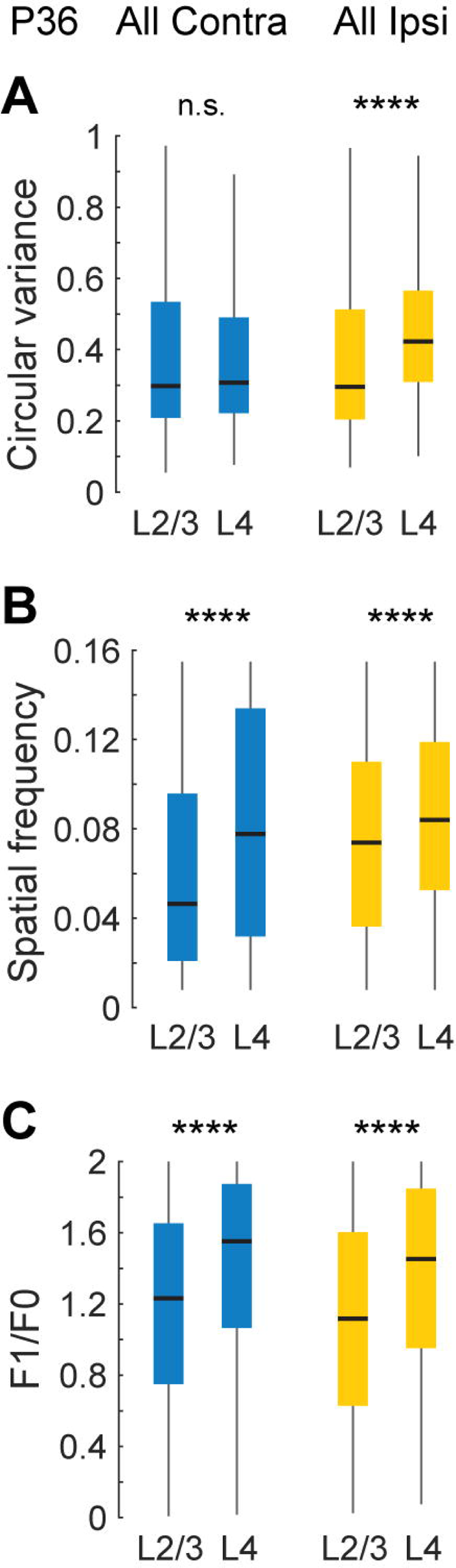
Receptive field tuning to ipsilateral eye in layer 2/3 is better than in layer 4 at P36 A. Boxplots of circular variance of neurons in layer 2/3 and 4 obtained via stimulation of the contralateral eye (blue, L2/3: 920 cells; L4: 673 cells) or ipsilateral eye (yellow, L2/3: 832 cells; L4: 848 cells). Mann-Whitney U test, ****, p<0.0001. B. As in A, but for spatial frequency preference. C. As in A, but for response linearity (F1/F0).

## Discussion

We quantify the development of receptive field tuning in layer 2/3 and 4 in mouse binocular V1. These measures complement existing studies that characterize the development of receptive field tuning properties in monocular primary visual cortex (Hoy and Niell, 2015). In agreement with others (Frégnac and Imbert, 1978; Freeman and Ohzawa, 1992; Crair et al., 1998; Issa et al., 1999; Smith and Trachtenberg, 2007; Jenks and Shepherd, 2020), we found few binocular neurons at eye opening. Instead, a majority of layer 2/3 pyramidal neurons in the binocular region respond solely to contralateral eye stimulation at this age. Consistent with earlier studies, these monocular, contralateral eye driven responses are selective across a range of receptive field tuning properties and are adult like in their preferences for spatial frequency and in the linearity of their responses (Hoy and Niell, 2015). Orientation selectivity of these neurons, which is poor at eye opening, becomes adult like by P18 (Li et al., 2012).

Ipsilateral eye-evoked responses are infrequent at eye opening but emerge quickly thereafter. Initially, receptive field tuning measures of ipsilateral-eye evoked responses are quite a bit worse than those from contralateral-eye evoked responses. Ipsilateral-evoked tuning improves exponentially after eye opening and becomes as good as contralateral-eye evoked tuning by P29. An orientation tuning bias towards the cardinal axes is seen at all ages, but for monocular neurons this bias becomes progressively more prevalent with age. This bias in adult mice has been previously reported (Salinas et al., 2017; Scholl et al., 2017b).

Few binocular neurons are present at eye opening in layer 2/3 and those that are present have poorly selective receptive field tuning and have poor binocular matching. By P18 the pool of binocular neurons has expanded considerably and approaches that seen in adult mice. Receptive field tuning of these P18 binocular neurons, however, remains significantly worse than in adult mice. Tuning properties improve exponentially from eye opening until P29 when they become significantly better than the tuning properties of monocular neurons. Moreover, while poor at eye opening, binocular matching was near adult like by P18, though the range of differences in orientation preference was quite a bit larger at P18 than at P36. There was little change in binocular matching from P22 through P36. We note that other studies report poorer binocular matching (Wang et al., 2010; Sarnaik et al., 2014). Part of this discrepancy may be due to methodology. In particular, the use of anesthetized vs. alert mice and the use of gratings that are more finely spaced in orientation (every 10 degrees vs. 30 degrees) are potential causes. All studies, ours included, define orientation tuning preference as the center of mass of the orientation response distribution. In addition to the progressive improvements in binocular matching and receptive field tuning, we found that binocular neurons in young mice display an orientation tuning bias towards the cardinal axes, but this bias is progressively lost with age as binocular neurons become responsive to the full range of orientations. This is quite the opposite of what we found for monocular neurons. This improvement in binocular tuning, matching, and orientation representation may be needed for efficient foraging (Butler, 1973; Hoy et al., 2016; Han et al., 2017; Shang et al., 2019; Zhao et al., 2019; Johnson et al., 2021).

In layer 4, the most notable feature was that ipsilateral eye-evoked tuning was substantially poorer than what is seen in layer 2/3. Binocular tuning was, therefore, poorer as well. Moreover, layer 4 was decidedly more monocular than layer 2/3. The continuous refinement of layer 2/3 ipsilateral eye responses, therefore, is unlikely to be driven by antecedent improvements in layer 4. That is, this improvement appears to be native to layer 2/3. There is growing evidence that receptive field tuning properties in visual cortex are established de-novo from layer to layer. For example, direction selectivity in layer 2/3 appears to be unrelated to the selectivity of its inputs (Rossi et al., 2020), and, in layer 4, the selective tuning of thalamocortical inputs (Sun et al., 2015), appears to be disregarded and tuning established de-novo (Lien and Scanziani, 2018). This laminar independence has previously been reported in studies of ocular dominance plasticity and normal binocular development in primary visual cortex (Trachtenberg et al., 2000; Trachtenberg and Stryker, 2001), but see also (Frantz et al., 2020). Taken together, these data suggest that cell types in layer 2/3 may be more sensitive to, and instructed by, vision than those in other layers.

The data we present provide a comprehensive overview of the development of binocular receptive field tuning in layers 2/3 and 4 in mouse primary visual cortex. The binocular visual field in mice is increasingly studied as a model system for revealing behaviorally relevant neural circuitry. Binocular neurons are tuned for disparity (Scholl et al., 2013, 2017a; La Chioma et al., 2019, 2020) and this information on depth underlies visually guided predation (Butler, 1973; Hoy et al., 2016; Han et al., 2017; Shang et al., 2019; Zhao et al., 2019; Johnson et al., 2021). The data we report here may prove useful for future studies on the development of the circuitry guiding this behavior and the role of early vision in establishing this circuitry.

## Author Contributions

Conceptualization, L.T., J.T.T.; Methodology, L.T., J.T.T., and D.L.R.; Investigation and Analysis, L.T.; Writing – Original Draft, J.T.T.; Writing – Review & Editing, L.T., J.T.T. Funding Acquisition, J.T.T.; Resources, J.T.T.; Supervision, J.T.T.

## Acknowledgements

This study was funded by NIH R01EY023871 (J.T.T.).

---

**Figure 2-1. Proportions of neurons per field of view or per mouse** A. Proportions of neurons in each field of view (FOV) at each age that are unresponsive to our visual stimuli, respond solely to stimulation of the contralateral eye (Contra), respond solely to stimulation of the ipsilateral eye (Ipsi), or respond to stimulation of either eye (Bino). As in A, but for proportions of neurons in each mouse.

